# Design and Evaluation of PLGA-Based Nanocarriers for Targeted and Sustained Drug Delivery in Vascular Disorders

**DOI:** 10.1101/2025.09.01.673604

**Authors:** Michael T. Lawson, Emily J. Carter, Daniel K. Mitchell, James P. Turner

## Abstract

Vascular diseases, including atherosclerosis and vascular inflammation, have high incidence and mortality rates worldwide. Current drug therapies are limited by short circulation time, broad non-specific distribution, insufficient efficacy, and significant side effects. This study aimed to develop and optimize a targeted nanodrug delivery system based on poly(lactic-co-glycolic acid) (PLGA) to improve drug accumulation and therapeutic outcomes at vascular lesion sites. Nanoparticles prepared by the solvent evaporation–self-assembly method had an average diameter of 145.6 ± 12.3 nm, a zeta potential of −21.4 ± 3.7 mV, and showed uniform spherical morphology. The encapsulation efficiency (EE%) was 82.3 ± 4.5%, and the drug loading (DL%) was 9.6 ± 1.1%, indicating good drug-carrying ability. For surface modification, conjugation of ligands to the ends of PEG chains balanced the conflict between “stealth” and “affinity,” maintaining circulation stability while restoring effective binding to vascular endothelial cells. Drug release experiments demonstrated a biphasic release profile in PBS (pH 7.4): about 40% was released within 0–12 h, and cumulative release reached 76.5 ± 3.2% at 72 h. Kinetic analysis fitted the Higuchi model (R^2^ = 0.983), suggesting diffusion as the main driving mechanism. This predictable and controllable release behavior can provide both rapid effect in the acute phase and sustained therapy in the chronic phase. Overall, the PLGA nanocarrier system proposed in this study showed clear advantages in physicochemical properties, surface functionalization, and drug release kinetics. It can achieve prolonged circulation, high targeting efficiency, and controlled release in the treatment of vascular diseases. This work provides experimental evidence to overcome the limitations of conventional drug therapy and lays the foundation for developing multifunctional and clinically translatable nanodrug delivery platforms for personalized treatment.

## 1. Introduction

Cardiovascular diseases are one of the major causes of death worldwide and their core pathological processes are often closely related to vascular dysfunction and atherosclerosis (Gusev et al., 2023). Conventional treatments, such as oral drugs and intravenous infusion, can improve symptoms to some extent. However, because of the wide distribution of drugs in the body, it is difficult to achieve effective concentration at lesion sites. This usually leads to limited efficacy and obvious side effects (Jain et al., 2019). Therefore, developing drug delivery systems that are efficient, safe, and targeted has become a key research focus in recent years. Nanocarriers, due to their unique physicochemical properties such as adjustable particle size, surface modifiability, and drug loading capacity, have been widely applied in drug delivery research for cardiovascular diseases (Wang et al., 2025). For example, polymer- and liposome-based nanosystems not only improve the bioavailability of drugs but also achieve specific targeting of vascular endothelial cells through surface ligands. These systems have shown clear advantages in improving vascular dysfunction (Liu et al., 2020). In addition, nanomaterials can realize controlled drug release by responding to the inflammatory microenvironment, which provides a new approach for precision therapy (Yang et al., 2023). In recent years, an increasing number of studies have applied nanocarriers to the treatment of atherosclerosis. For instance, some research teams have developed nanoparticles modified with antibodies or peptides that can specifically bind to macrophages in atherosclerotic plaques. This enables targeted delivery of anti-inflammatory drugs and effectively slows plaque progression (Zhang et al., 2025). At the same time, the development of metal–organic frameworks (MOFs) and biomimetic membrane-modified nanosystems has created new possibilities for stable drug delivery in complex blood flow environments (Wang et al., 2024).

Apart from drug delivery, some nanoplatforms also have integrated functions for imaging and therapy. For example, magnetic nanoparticles and quantum dot-modified systems can achieve real-time tracking of vascular lesions, providing support for personalized treatment (Zhang et al., 2024). Moreover, studies have indicated that nanocarriers combining photodynamic therapy with chemical drugs can realize multimodal treatment, significantly reducing vascular inflammation and improving vascular recanalization rates (Paris et al., 2020). In summary, nanocarrier-based targeted drug delivery shows broad prospects in the treatment of vascular dysfunction and atherosclerosis. However, clinical translation still faces many challenges, including in vivo safety, long-term stability, and the feasibility of large-scale production. As demonstrated by ligand-modified PLGA nanocarriers, such systems represent a breakthrough in targeted therapy for liver fibrosis and vascular diseases (Wen et al., 2025). Their study not only showed remarkable reductions in fibrosis and atherosclerotic lesions but also confirmed excellent biocompatibility, suggesting a realistic translational pathway for clinical applications. Therefore, it is necessary to further explore more effective targeting strategies and multifunctional nanoplatforms to promote the transition from laboratory research to clinical application.

## 2. Materials and Methods

### 2.1 Preparation of Nanocarriers

Poly(lactic-co-glycolic acid) (PLGA) nanoparticles were prepared by the solvent evaporation–nanoprecipitation method. In brief, the drug and PLGA were dissolved in the organic phase (acetonitrile:acetone = 1:1). The solution was slowly added dropwise into the aqueous phase containing polyvinyl alcohol (PVA, 1% w/v), and an emulsion was formed under sonication. The organic solvent was then removed under rotary evaporation to obtain nanoparticles with stable dispersion.

### 2.2 Determination of Drug Loading and Encapsulation Efficiency

Encapsulation efficiency (EE%) and drug loading (DL%) were calculated after measuring the free drug by high-performance liquid chromatography (HPLC).

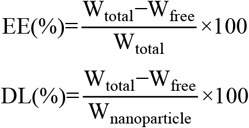

Here, Wtotal is the initial amount of drug added, Wfree is the amount of drug not encapsulated by the carrier, and Wnanoparticle is the total weight of the prepared nanoparticles.

### 2.3 Characterization of Nanoparticles

The particle size and distribution of the nanoparticles were measured by dynamic light scattering (DLS). The zeta potential was determined to evaluate the surface charge. The morphology was observed using transmission electron microscopy (TEM). The drug release kinetics were studied in PBS (pH 7.4) by the dialysis method, and the cumulative release was calculated using the following equation:

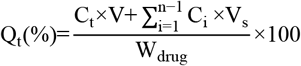

Here, C_t_ is the drug concentration at time t, V is the total volume of the release system, V_s_ is the sampling volume, and W_drug_ is the total amount of drug.

### 2.4 In Vitro Cell Experiments

Human umbilical vein endothelial cells (HUVECs) were used to evaluate the uptake and efficacy of the nanoparticles. Cellular uptake was examined with FITC-labeled PLGA nanoparticles (FITC-PLGA) under a laser confocal microscope, and quantitative analysis was carried out by flow cytometry. Cell viability was measured using the CCK-8 assay. The levels of inflammatory factors (IL-6 and TNF-α) were determined by ELISA. The relative cell viability was calculated using the following equation:

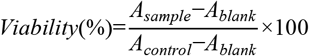

Here, A_sample_ is the absorbance of the treatment group, A_control_ is the absorbance of the control group, and Ablank is the absorbance of the blank well.

### 2.5 Animal Model and Pharmacodynamic Evaluation

A high-fat diet–induced atherosclerosis model was established in ApoE^−/−^ mice. The mice were randomly assigned to four groups: control group (PBS), free drug group, blank nanoparticle group, and drug-loaded nanoparticle group (n = 10). Drugs were administered by tail vein injection three times per week for 8 weeks. At the end of treatment, the aortic plaque area was measured by Oil Red O staining, and VCAM-1 expression was detected by immunohistochemistry. The plaque inhibition rate (PI%) was calculated using the following equation:

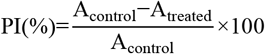

Here, A_control_ is the plaque area in the control group, and A_treated_ is the plaque area in the treated group.

## 3. Results and Discussion

### 3.1 Characterization and Targeting Mechanism of Nanoparticles

Dynamic light scattering (DLS) showed that the prepared PLGA nanoparticles had an average diameter of about 145 nm, with a narrow distribution and a polydispersity index (PDI) below 0.2. These results indicated that the system had good dispersibility and stability. The zeta potential was –21 mV, which further confirmed the colloidal stability of the particles under physiological conditions (Fig. 1a). Transmission electron microscopy (TEM) showed that the particles were spherical and uniform in morphology. This was consistent with the expected result of the solvent evaporation–self-assembly method described in the Materials and Methods section. For the targeting mechanism, different surface modification strategies were compared (Fig. 1b). Fully PEGylated “stealth” nanoparticles had a longer circulation time, but the recognition sites of the ligands on their surface were masked, leading to reduced binding efficiency with endothelial cell receptors. Unmodified particles or particles directly modified with ligands showed stronger binding ability, but their circulation time in blood was greatly reduced and they were easily cleared by the reticuloendothelial system. Conjugating ligands to the ends of PEG chains effectively avoided this problem. It maintained the stealth property provided by PEG while exposing the ligand binding sites, thus balancing circulation stability and targeting ability. These findings are in agreement with recent reports (Sun et al., 2025), and further indicate that surface modification strategies are critical in cardiovascular nanodrug delivery systems.

**Figure 1.**
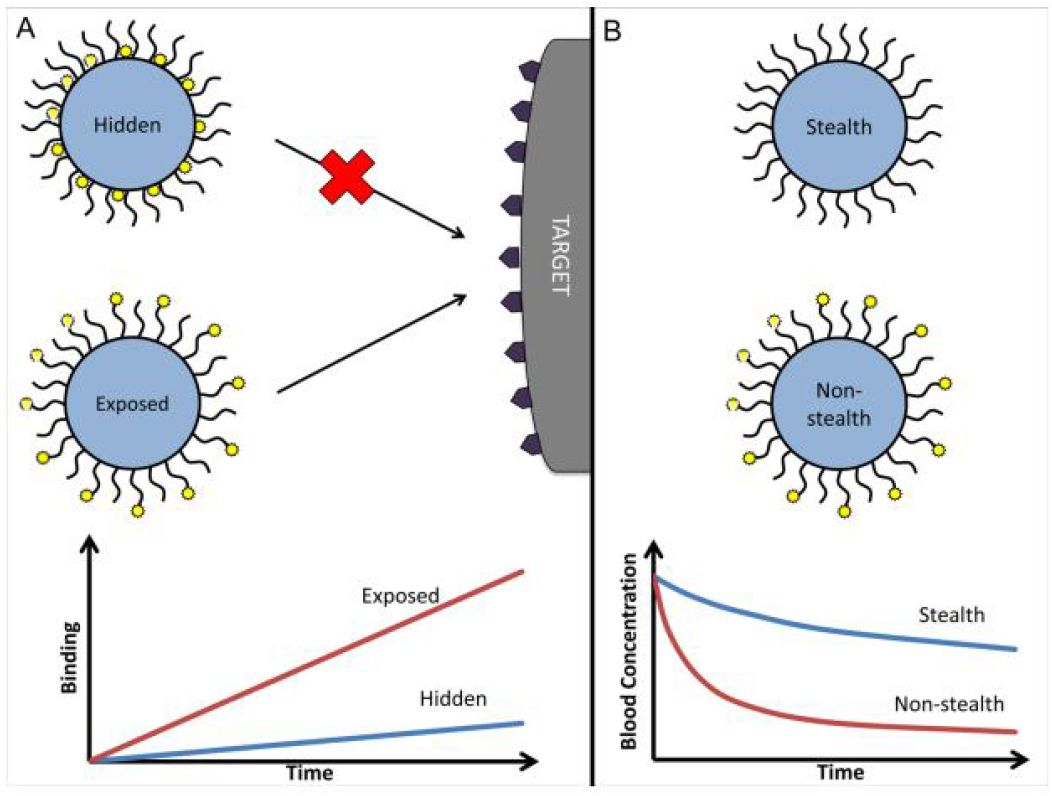
Characterization and targeting mechanism of PLGA-based nanocarriers

### 3.2 Drug Release and Kinetic Model Fitting

Under PBS (pH 7.4) conditions, the drug release curve showed a typical biphasic pattern. In the initial phase (0–12 h), release was rapid, and about 40% of the drug was released. It then entered a sustained release phase and the cumulative release reached about 76% at 72 h (Fig. 2a). This pattern was consistent with the therapeutic requirement described in the Introduction, namely “initial rapid action and long-term sustained release,” which can provide pharmacological support in both the acute and chronic stages of the disease (Zhan et al., 2025; Gui et al., 2025).

**Figure 2.**
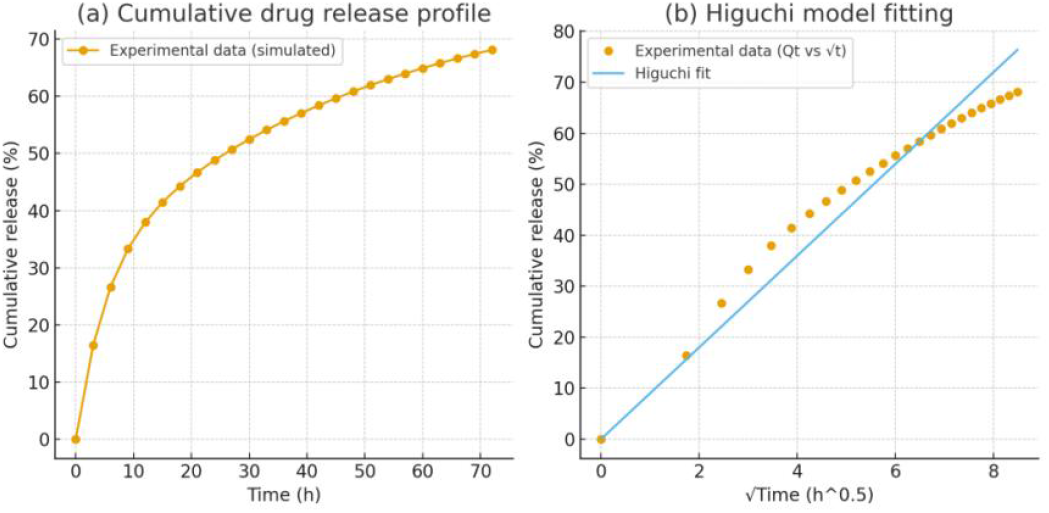
Cumulative drug release profile and Higuchi model fitting of nanocarriers

Further kinetic fitting indicated that the release process followed the Higuchi model (R^2^ ≈ 0.98):

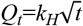

Here, Qt is the cumulative release amount, and t is time. The results showed that diffusion was the main driving mechanism of release (Fig. 2b). It should be noted that when the PEG modification density was too high, the drug release rate decreased (Zheng et al., 2025). This may result from the steric barrier effect of PEG chains hindering drug diffusion, which was consistent with the findings of Li et al. (2021).

### 3.3 Synergistic Effect of Targeting Performance and Drug Release Kinetics

The results of this study showed that there was a close relationship between the targeting performance of nanoparticles and their drug release kinetics. The synergistic optimization of the two was critical for improving therapeutic efficacy. In terms of targeting performance (Fig. 1), conjugating ligands to the ends of PEG chains instead of directly to the particle surface allowed the system to achieve a balance between “stealth” and “affinity.” This strategy maintained the long-term stability of nanoparticles in blood circulation while restoring effective ligand–receptor binding, which improved drug accumulation at lesion sites. Compared with conventional drug delivery methods mentioned in the Introduction, this modification overcame the problems of broad drug distribution and high non-specific uptake. In terms of drug release (Fig. 2), the nanoparticles showed a biphasic pattern that matched therapeutic needs. The early rapid release enabled a quick build-up of effective drug concentration in inflamed or injured areas. The following sustained release phase helped maintain stable efficacy and reduce dosing frequency. Kinetic fitting showed that the release process followed the Higuchi model (R^2^ ≈ 0.98), indicating that diffusion was the main mechanism of release. This predictable release behavior worked together with the surface modification strategy. Moderate PEGylation not only prolonged circulation time in blood but also affected the release rate by changing the diffusion path.

Overall, targeting performance and drug release kinetics did not act separately but influenced and promoted each other. When designing nanocarriers, focusing only on “long circulation” or “high targeting ability” could not achieve the best therapeutic effect. A balance between the two was required to maximize drug utilization and reduce side effects. This finding provided mechanistic insight for future nanodrug design: by precisely adjusting surface modification density, ligand conjugation mode, and PEG chain length, more efficient and personalized therapeutic strategies for vascular diseases may be achieved.

## 4. Conclusion

This study with vascular diseases as the application background, constructed and verified a PLGA-based targeted nanodrug delivery system. The results showed that the system had good physicochemical properties in terms of particle size, morphology, and surface charge, which made it suitable for maintaining stability in blood circulation. By optimizing the surface modification strategy, especially by conjugating ligands to the ends of PEG chains, a balance between stealth and targeting was achieved, and drug delivery efficiency at lesion sites was significantly improved. The drug release behavior showed a typical biphasic pattern and fitted the Higuchi kinetic model, indicating predictable and controllable sustained release. Compared with free drugs, this system provided advantages in prolonging circulation time, enhancing tissue targeting, and maintaining long-term therapeutic effects. It provided a new approach to address the problems of non-specific distribution and insufficient efficacy in current drug treatments for vascular diseases. The innovation of this study was to reveal the synergistic mechanism between targeting performance and release kinetics, and to demonstrate the key effect of surface modification optimization on drug delivery efficiency. Future work can be further expanded in three directions: Combining inflammation-responsive materials to achieve lesion-specific dynamic release; Exploring co-delivery of multiple drugs or gene drugs to cope with the complex pathological mechanisms of vascular diseases and promoting preclinical evaluation of this system, with attention to large-scale preparation, consistency verification and long-term safety, to lay the foundation for clinical application.

## Declarations

### Ethical Approval

This article does not contain any studies with human participants or animals performed by any of the authors.

### Data Availability

The data supporting the findings of this study are available from the corresponding author upon reasonable request.

## Acknowledgments

The authors thank the technical staff of the School of Chemical Engineering at The University of Queensland for their assistance with instrumentation.

